# Cassava Detection from UAV Images Using YOLOv5 Object Detection Model: Towards Weed Control in a Cassava Farm

**DOI:** 10.1101/2022.11.16.516748

**Authors:** Emmanuel C. Nnadozie, Ogechukwu Iloanusi, Ozoemena Ani, Kang Yu

## Abstract

Most deep learning-based weed detection methods either yield high accuracy, but are slow for real-time applications or too computationally intensive for implementation on smaller devices usable on resource-constrained platforms like UAVs; on the other hand, most of the faster methods lack good accuracy. In this work, two versions of the deep learning-based YOLOv5 object detection model – YOLOv5n and YOLOv5s - were evaluated for cassava detection as a step towards real-time weed detection. The performance of the models were compared when trained with different image resolutions. The robustness of the models were also evaluated under varying field conditions like illumination, weed density, and crop growth stages. YOLOv5s showed the best accuracy whereas YOLOv5n had the best inference speed. For similar image resolutions, YOLOv5s performed better, however, training YOLOv5n with higher image resolutions could yield better performance than training YOLOv5s with lower image resolutions. Both models were robust to variations in field conditions. The speed vs accuracy plot highlighted a range of possible speed/accuracy trade-offs to guide real-time deployment of the object detection models for cassava detection.

## Introduction

There is currently a global threat to food security, which is attributed mainly to world population growth, conflicts, and climate change (Food and Agriculture Organisation of the United Nations 2017; United Nations 2019). Consequently, there is an increasing call for research directed at developing sustainable solutions to the food challenge; such that agrochemical inputs are minimized while increasing yield. These challenges present opportunities for innovations in farming technologies; thus encouraging the adaptation of existing cutting-edge technologies and the development of new methods (Duckett et al. 2018).

Weed detection is an important step in weed management. Not only is weed detection important to the farmers for weed control tasks, but recently, ecologists have begun to highlight the importance of weed detection for biodiversity monitoring. For instance, in (Adair and Richard 1998; Balasubramanian et al. 2014) weed detection is highlighted as an essential step in determining the ecological distribution of weeds, understanding their impact on the environment, as well as establishing an information system for their management. Economic, environmental, as well as health concerns constitute the challenges facing the prevalent weed control approach using herbicides; especially because the current practice involves broadcast spraying of the entire field with the same dosage (Devos et al. 2008; Fillols et al. 2020; Liu and Bruch 2020). Thus, the need for solutions for per plant treatment is emphasized.

Manual inspection of farms to identify weeds can be labourious and time-consuming. Therefore, the development and deployment of autonomous and semi-autonomous farm vehicles including terrain vehicles and, more recently, unmanned aerial vehicles (UAVs) offer great potentials and alternative approaches for site-specific weed management. Such vehicles are fitted with perception systems based on various computer vision techniques for weed detection (Ukaegbu et al. 2021). Traditional image processing techniques such as colour thresholding, filtering, and differencing are methods first studied by researchers for weed detection (Mustafa et al. 2007). More recently, learning-based methods like Support Vector Machines, Artificial Neural Networks, Random Forests, among others (Barrero et al. 2016; Saha et al. 2016; Wu et al. 2021) have been shown to have superior performance over traditional image processing methods for weed detection. While traditional machine learning methods showed promising results in controlled environments, their accuracy is undermined in real-life field conditions. Their lack of robustness to field conditions like natural lighting, varying plant density, leaf occlusions, and different plant growth stages necessitated the development of deep learning methods for weed detection. In addition to the robustness of deep learning methods to field variations, they are capable of end-to-end learning of hierarchical features of plant images, thus eliminating the need for the labourious task of feature engineering required in machine learning.

Convolutional Neural Networks including AlexNet, VGG16, Inception-ResNet, etc. are among the deep learning methods employed for weed detection as demonstrated in (Bah et al. 2019; Espejo-Garcia et al. 2020; Tang et al. 2017). However, object detection, in contrast to whole image classification, is more useful for weed identification and localization in weed control. Region proposal-based models such as Faster-RCNN and Mask-RCNN were used by (Khan et al. 2021; Osorio et al. 2020; Thanh Le et al. 2021) for weed detection. While such models deliver promising results in terms of accuracy, because they use a two-stage detection pipeline, they have slower inference speeds than their regression-based counterparts like the Single Shot Multibox Detector (SSD) and YOLO networks (Huang et al. 2017; Osorio et al. 2020).

The YOLO object detection model was introduced by Redmon et al (Redmon et al. 2016). Unlike previous object detection models such as Faster R-CNN (Ren et al. 2017) which employ a two-stage process of region proposals followed by bounding box classification, YOLO is a single-stage detector performing all the object detection processes in a single network. The regression-based YOLO model overcame the bottlenecks posed by the complex architecture of the region-based detectors, greatly reducing the inference time, while enhancing generalizability of the model. Thus, the YOLO is favoured for real-time applications.

The first YOLO model was quickly followed by the release of YOLOv2 (Redmon and Farhadi 2017) which addressed limitations of the first version such as lack of generalization when image objects are too small. YOLOv3 (Redmon and Farhadi 2018) was released to ameliorate the accuracy issues encountered in YOLOv2. The release of YOLOv4 (Bochkovskiy et al. 2020) came with better accuracy, and was followed by YOLOv5 (Jocher et al. 2020) with a much lighter weight file. While YOLOv1 to YOLOv4 were implemented on the rather unpopular Darknet framework, YOLOv5 is implemented on the widely used and research-friendly Pytorch deep learning framework. Fig. 1 shows the performance of the various YOLOv5 models on the COCO dataset.

**Fig. 1.**
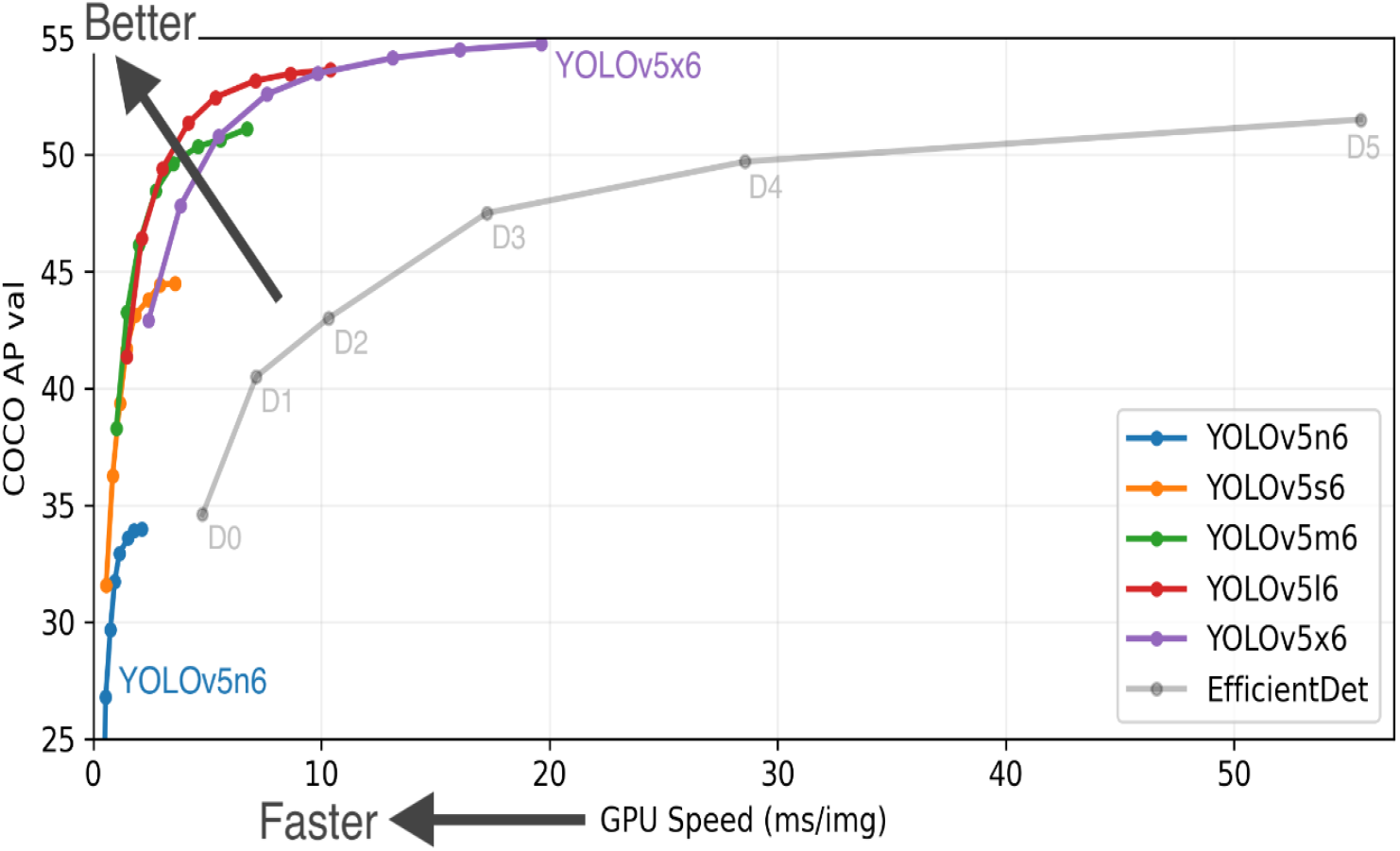
YOLOv5 performance on COCO dataset (Jocher et al. 2020)

YOLO networks have been deployed in weed detection for high inference speed which makes them attractive for real-time applications. YOLOv3 was deployed on a ground robot for real-time weed detection in a carrot farm; and the setup achieved up to 89% precision (Czymmek et al. 2019). Gao et al developed a detection model based on Tiny YOLO for detecting *Convolvulus sepium* in sugar beet fields, and achieved up to 0.897 accuracy (Gao et al. 2020).

In spite of the plethora of works on weed detection using deep learning methods, it is not common to find research on real-time weed detection using UAV imagery, despite the many advantages of UAVs; such as ability to operate irrespective of the terrain, and the elimination of unwanted soil impact and damage as is common with ground vehicles. Furthermore, much of the existing research in weed detection and control is focused on crops and weeds cultivated in the western countries, and not on crops peculiar to sub-Saharan Africa such as cassava *(Manihot esculenta)* which is a major food crop in sub-Saharan Africa (Food and Agriculture Organisation of the United Nations 2010).

In this paper, the focus is on real-time cassava detection from UAV images as a first step to weed detection and control on a cassava farm in Nigeria using unmanned aerial vehicles. The performance of YOLOv5n and YOLOv5s (Jocher et al. 2020) object detection models on our dataset were evaluate and compared. Also, the performance of the models on different input image resolutions were evaluated in order to highlight the potential speed/accuracy trade-offs based on model type and image size. Furthermore, evaluation and comparison of the inference performance of the models were carried out for different field scenarios including varying lighting and shadow presence, different cassava growth stages, varying weed density and plant leaf occlusions, as well as different row orientations.

Our contributions were as follows:

1. Creation of a dataset of UAV cassava images for weed detection.
2. Implementation of YOLOv5n and YOLOv5s for real-time cassava detection.
3. Evaluation of YOLOv5n and YOLOv5s for cassava detection under varying growth stages, natural lighting, and weed density.
4. Assessment of the speed/accuracy trade-offs for cassava detection with respect to varying input image resolution and YOLOv5n and YOLOv5s model types.

## Materials and Methods

This section will describe the various techniques and materials employed for the realization of the work.

### Dataset

The performance of object detection models depends a lot on the quality of the data the model is trained on. Therefore, it was necessary to carefully compile a suitable dataset for the training of the YOLOv5 model.

#### Experimental setup

For this research, an experimental cassava farm was set up some kilometers away from the University of Nigeria, Nsukka. The farm setup process included land selection, clearing, ploughing, ridging, planting, and manuring. The cassava cuttings were planted at distances of 1m apart according to the standard recommendations by International Institute of Tropical Agriculture (IITA) for cassava cultivation in the derived savanna region of Nigeria (Hauser et al. 2014).

#### Image Acquisition

A 550mm quadcopter was constructed for the purpose of capturing aerial images of the farm. A GoPro Hero 7 camera was mounted on the quadcopter to capture RGB images of the farm. The farm images were captured at altitudes ranging between 2 meters to 3 meters between the months of September and October, 2021, specifically, 25, 29, and 60 days from planting. The images were collected under different natural lighting conditions, cassava growth stages, as well as varying weed densities and leaf occlusions. Each of the captured images were of resolution 4000 x 3000 pixels.

#### Image annotation

One hundred images and 25 images were respectively used for training and validation of the models, while 18 images were used for testing. Only the train and validation sets were annotated. The test images were used for inference purposes and therefore did not require annotation. The annotation of the images involved drawing bounding boxes around the objects of interest and assigning object classes to the bounding boxes. The YOLOv5 requires that the bounding box dimensions be normalized to the range [0,1] and represented in the “c x y w h” format; where c is the object class index (starting from 0), x and y represent the x-y coordinates of the center of the bounding box, and w and h represent the width and height of the bounding boxes respectively. For each image, the annotations must be saved in a .txt file, where each bounding box information is entered on a new line. The python-based LabelImg (Tzutalin 2015) graphical annotation tool was chosen for image annotation in this work because of its ease of use and ability to save the annotations in the YOLOv5 format.

#### Data augmentation

To avoid overfitting to the data, this work defaulted to the use of the inbuilt train-time data augmentation of YOLOv5. This includes image-space and colour-space transformations and the novel mosaic data augmentation proposed by the author of YOLOv5. The mosaic augmentation performs random image augmentations of the original image and assembles it into a mosaic image consisting of the original image and a number of randomly augmented images. The data augmentation can be modified in the hyperparameters which includes options for specifying or even disabling augmentation types and values. By adjusting the augmentation hyperparameters, our model was exposed to broader semantic representations of the training data.

### Object detection

The following section will describe the YOLOv5 object detection model, the training platform.

#### Model Training

The YOLOv5 release comprised of YOLOv5n, YOLOv5s, YOLOv5m, YOLOv5l, and YOLOv5x variants with accordingly increasing weight file size, training time, number of parameters, and model accuracy. Thus, YOLOv5n has the smallest weight file, least training time and number of parameters, making it suitable for deployment on memory constrained devices. Conversely, YOLOv5x the largest and most accurate of the variants is best suited for applications where very high accuracy is a priority and memory is not a problem. Since our target was for real-time applications, in this work, the two smallest models, YOLOv5n and YOLOv5s, were chosen for their lightness, high inference speeds, and acceptable accuracy.

The YOLOv5 model training and inference was done on NVIDIA GeForce 3060 on our local workstation using Jupyter Notebooks.

The model was trained on 100 images and validated on 25 images. As a rule of thumb, the training data for object detection should contain about 1500 objects per class. Counting the number of objects in our training data using our python script gave us over 1700 objects for the cassava class, which was the only object class.

The model was trained for different combinations of input image and batch sizes. These combinations were also trained on multiple models of the YOLOv5 as mentioned previously. The range of image input sizes include 960×720, 640×480, 512×384, and 256×192; while the batch size ranged from 8 to 64. The maximum batch sizes for training on 960×720 and 640×480 image sizes were limited to not more than 20 and 32 respectively, because the GPU memory did not permit loading more than these limits for the respective image sizes. The results were evaluated to identify which model and parameter combinations gave the best performance for real-time applications. For all the training instances, a fixed epoch of 400 was used.

Given that for computer vision tasks, using pretrained weights can lead to better accuracy than using random weights (Hertel et al. 2015), the pretrained weights corresponding to the respective models, which were trained on the COCO dataset (Lin et al. 2015) by the YOLOv5 author, were used. Initial learning rate at 0.01 and momentum of 0.937 was used, while the weight decay was set at 0.0005.

#### Performance metrics

The precision and recall, and mean average precision of the model were used for evaluation of the model performance. The precision is the degree to which the model correctly predicts the objects. The precision, P is represented by the relation

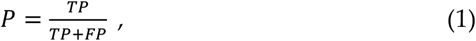

Where TP is true positives, FP is false positives. As observed from equation (1), the higher the false predictions, the lower the precision.

Recall, on the other hand, is the degree to which the model captures all ground truth. Recall, R is given by equation (2)

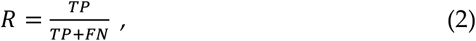

Where FN is false negative.

The mean average precision, mAP, has become the acceptable metric in the research community for evaluating object detection models. The mAP indicates the weighted mean of precisions obtained for different intersection over union (IOU) thresholds. In this work, the precision, recall, mAP@0.5, and mAP@0.5:0.95 of all models were extrapolated.

## Results and Discussion

### Captured UAV images

Fig. 2 shows sample images captured under various field conditions.

**Fig. 2.**
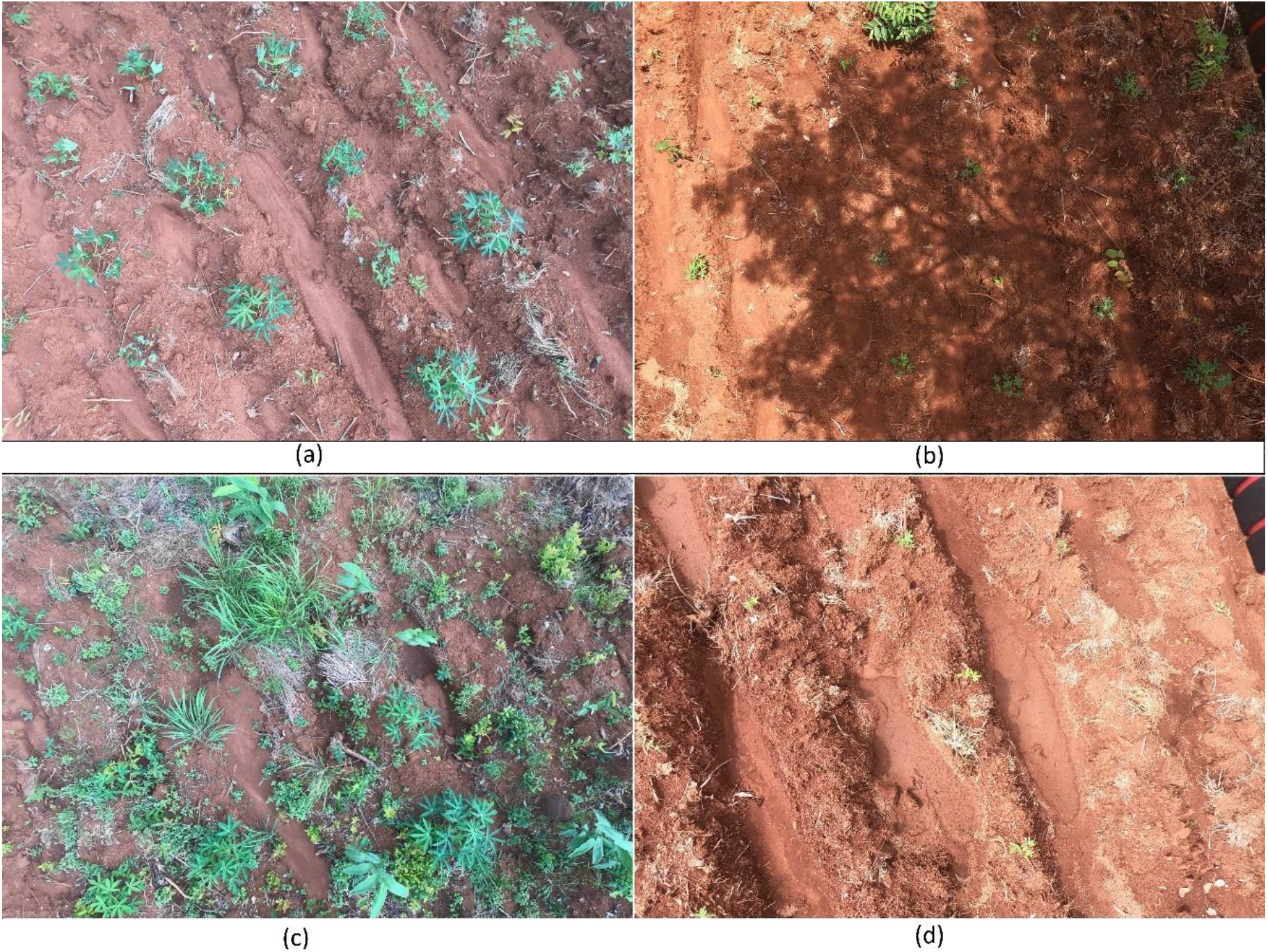
UAV Field images showing varying field conditions; (a) low lighting, low weed density, and vegetative cassava growth stage, (b) high illumination with shadow, (c) high weed density with high leaf occlusion, (d) high illumination with early cassava growth stage

To improve the robustness of the models, images were captured in different row orientations. Images showing different row orientations are shown in Fig. 3.

**Fig. 3.**
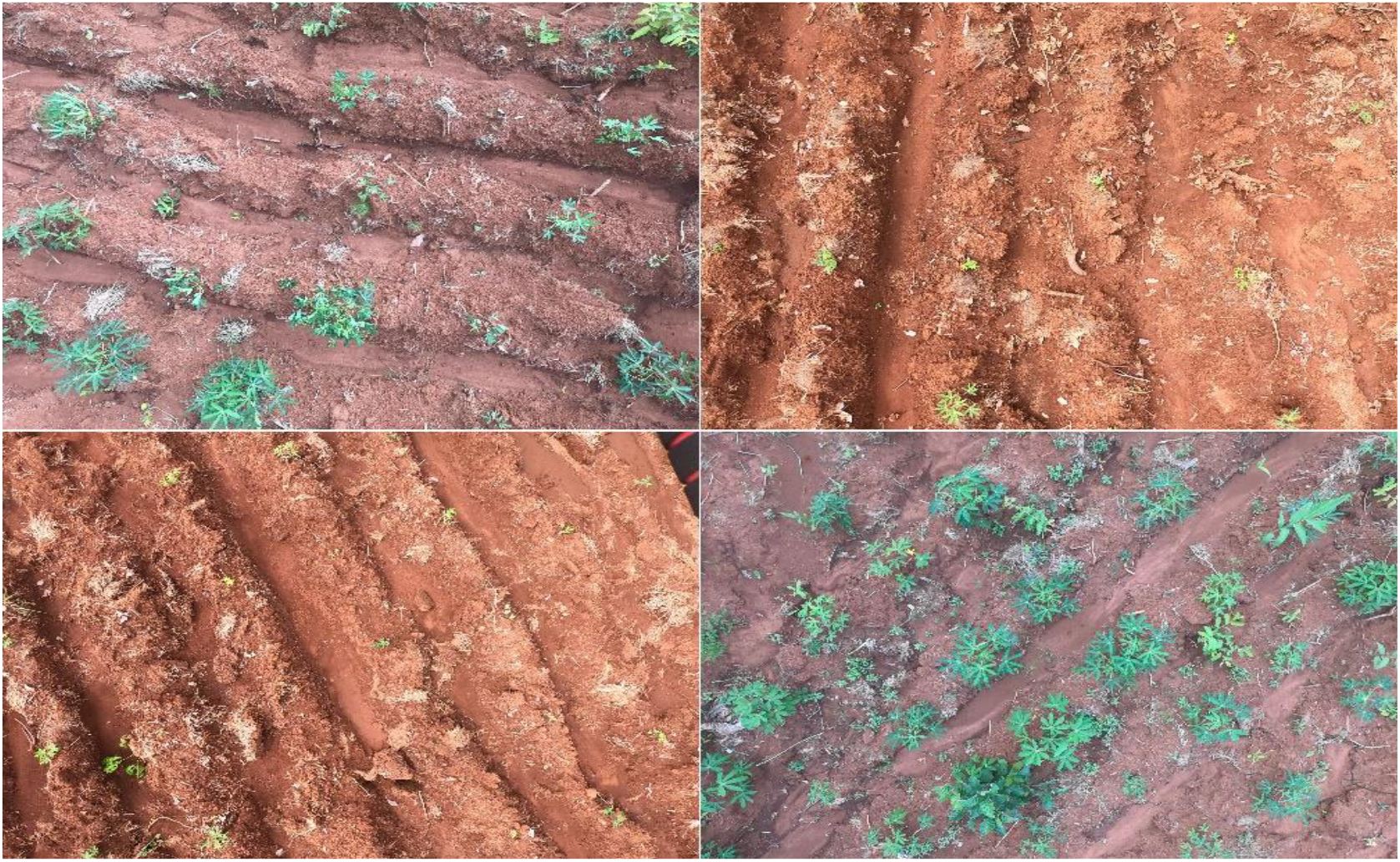
UAV field images showing different row orientations

Some annotated images are shown in Fig. 4. The bounding boxes are shown to be drawn as tightly as possible about the cassava plants to enhance learning of the cassava features by the models. The dataset together with the annotation files are available upon request from the authors.

**Fig. 4.**
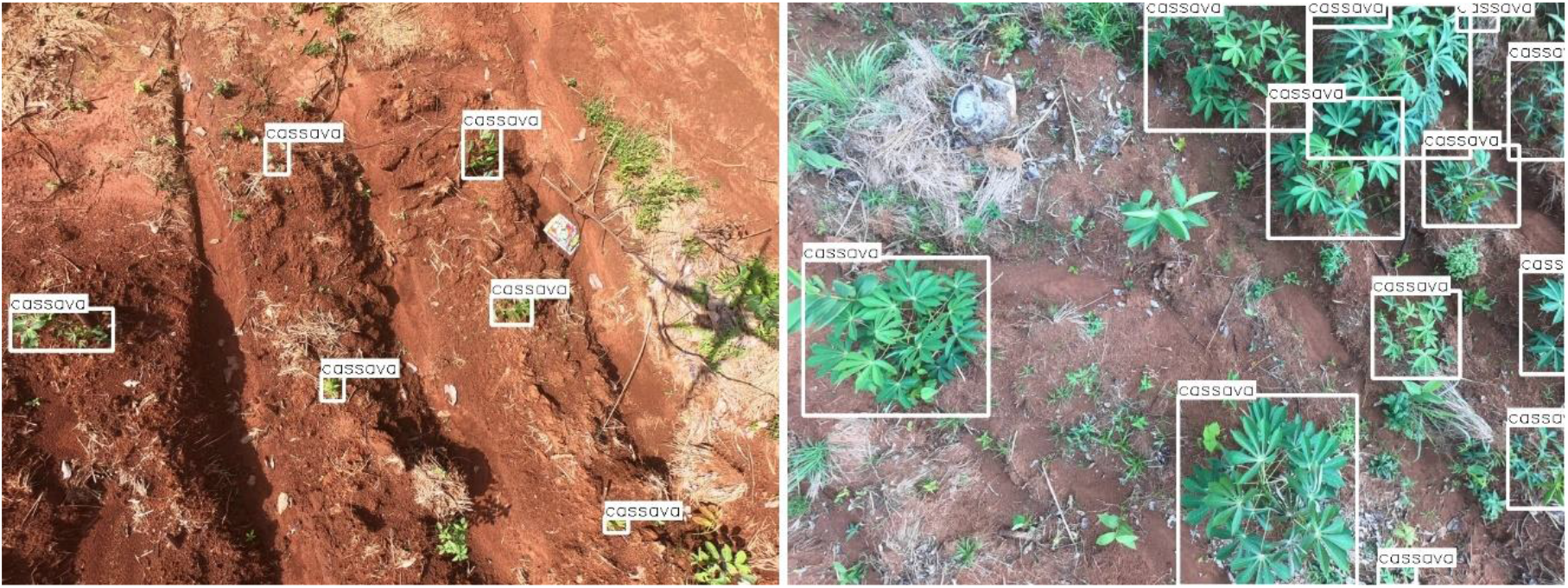
Annotated field images with object class labels

### Model Training Performance

Table 1 shows the summary of the training results. The training duration, precision, recall, mAP@0.5, and mAP@0.5:0.95 are shown for different input image sizes, batch sizes, as well as for the two YOLOv5 models. While the aforementioned performance metrics are presented in the table, the discussion will consistently reference the mAP@0.5:0.95 which is the more widely accepted performance metric in the object detection research community.

**Table 1.**
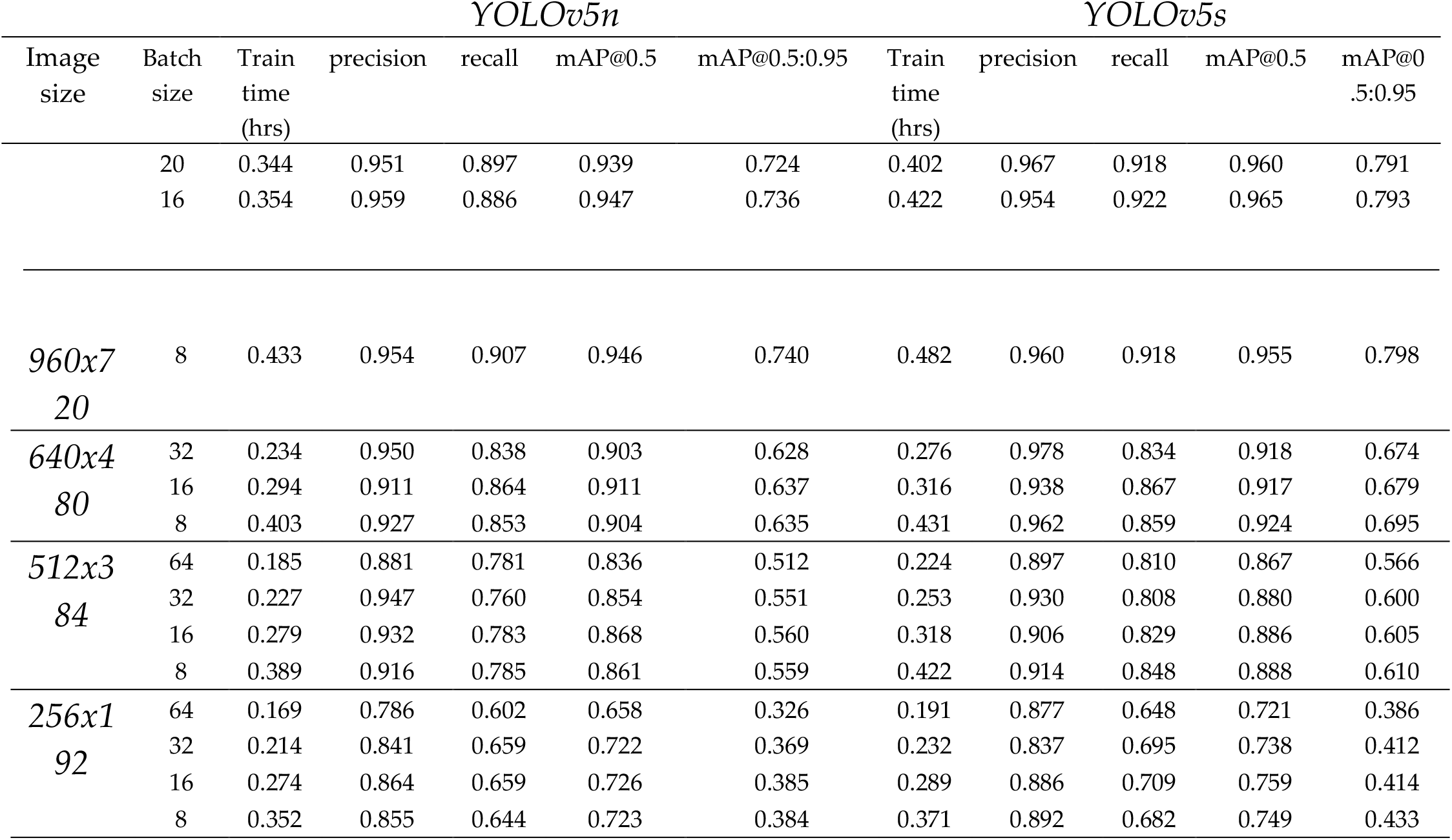
Summary of training results

#### Performance of the models for varying image sizes

Fig. 5 shows the curves of the best mAP@0.5:0.95 for each image resolution for each YOLOv5 model.

**Fig. 5.**
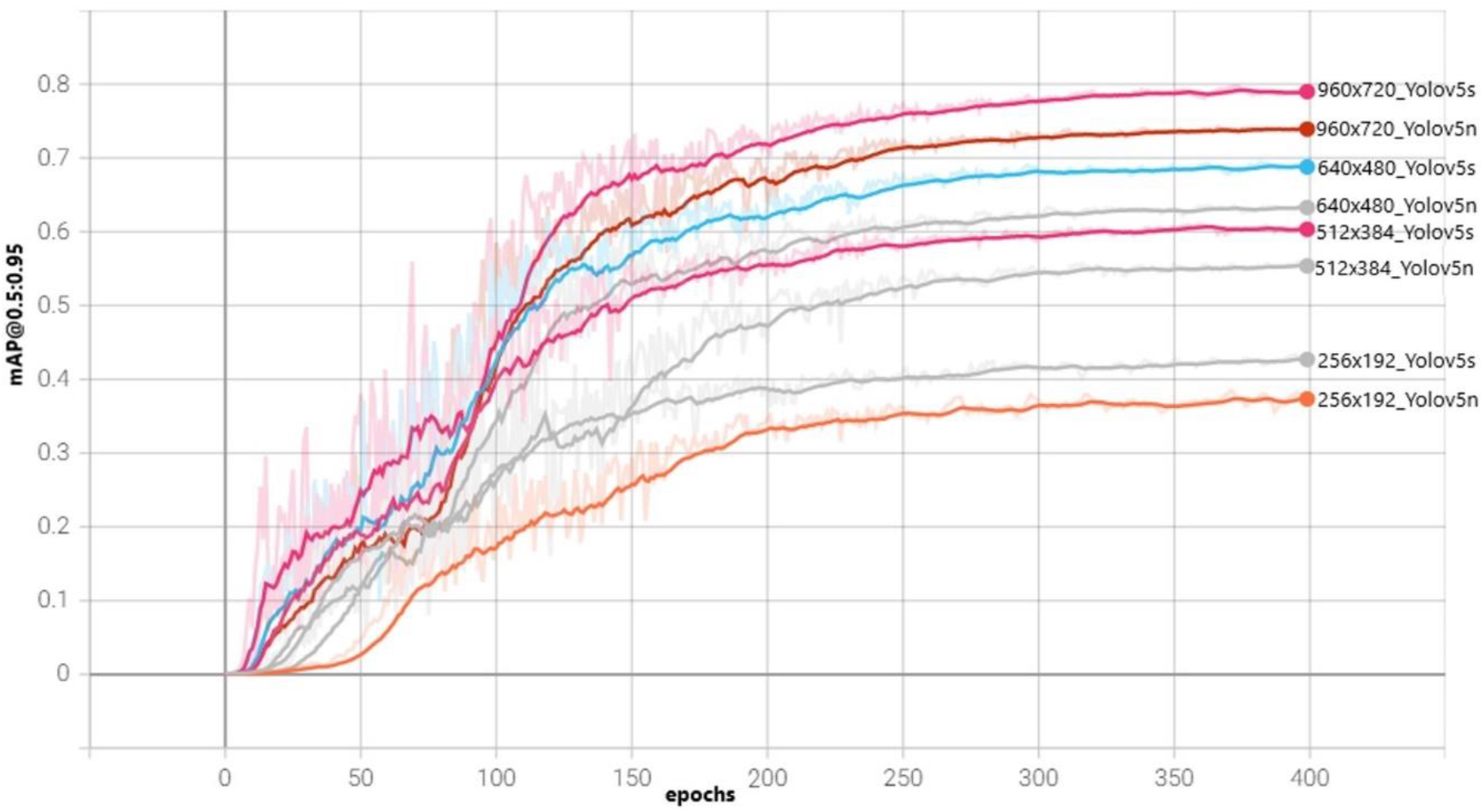
mAP@0.5:0.95 for various image resolutions

The mAP@0.5:0.95 decreases with decrease in image resolution. This decrease in performance is because it is more difficult for the models to resolve small objects as the image resolution decreases. However, the increase in performance with increase in image resolution comes at the cost of increased training time for the models, as shown in the chart in Fig. 6. This is due to the fact that the higher the image size, the more memory and computational resources are required to process it; as well as more features to learn. Therefore, for applications where training time is critical or memory constraint is high, lower image resolutions may be used, but at the expense of model accuracy.

**Fig. 6.**
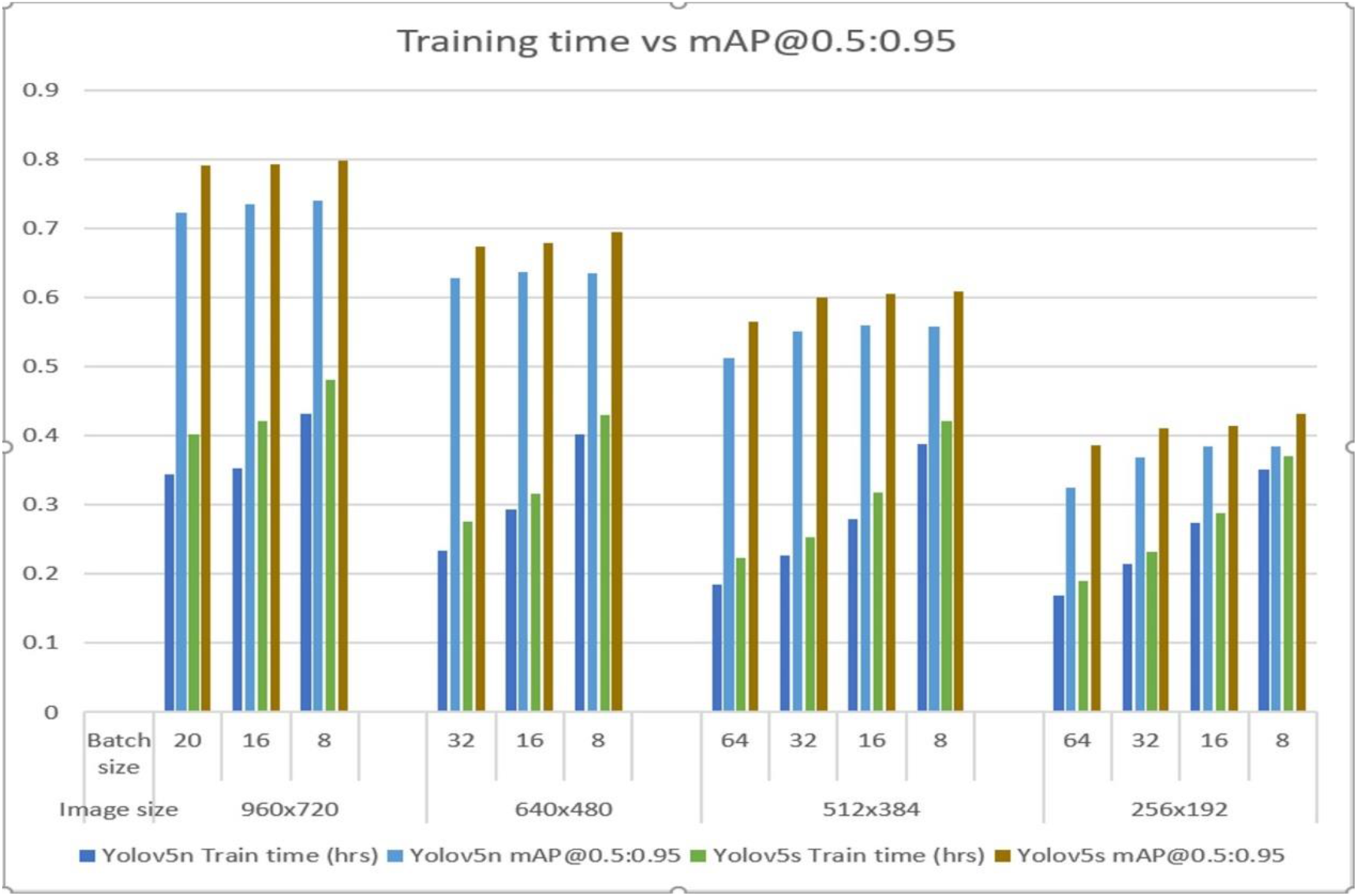
Training time and mAP@0.5:0.95 for all image resolutions and batch sizes

It is also noted that for each image resolution, the YOLOv5s model performs better than the YOLOv5n model. The better performance of YOLOv5s is because it is able to learn better with more parameters than the YOLOv5n. But this is at the expense of having a heavier weights file than the YOLOv5n. For instance, the YOLOv5s learns with over 7 million parameters and has a weight file of 14.6Mb, whereas the YOLOv5n has less than 2 million parameters with a weight file size three times less at 4.1Mb.

Interestingly, as observed in the chart on Fig. 6, an interplay of batch size, image size, and model selection could lead to better performance of the YOLOv5n model. For instance, it is extrapolated from our results, as shown in Fig. 6, that training YOLOv5n on 960×720 images yields higher mAP@0.5:0.95 than training YOLOv5s on any of the lower image resolutions. The only cost is a slightly higher training time. This may be desirable for applications such as realtime weed detection from UAV images where the resolution needs to be as high as possible and the detection model as small as possible to fit the memory constraints for UAV-mounted processors.

#### Effect of varying batch sizes on the model performance

For all image sizes, the training time increases with decrease in batch size as shown in Fig. 6. This implies that taking more images per batch is more memory efficient. However, from the mAP@0.5:0.95 curve for different batch sizes of 960×720 input image size shown in Fig. 7, which is fairly representative of other image sizes, the difference in mAP@0.5:0.95 is not very significant. Thus, we conclude that the major advantage of higher batch sizes is the decrease in training time.

**Fig. 7.**
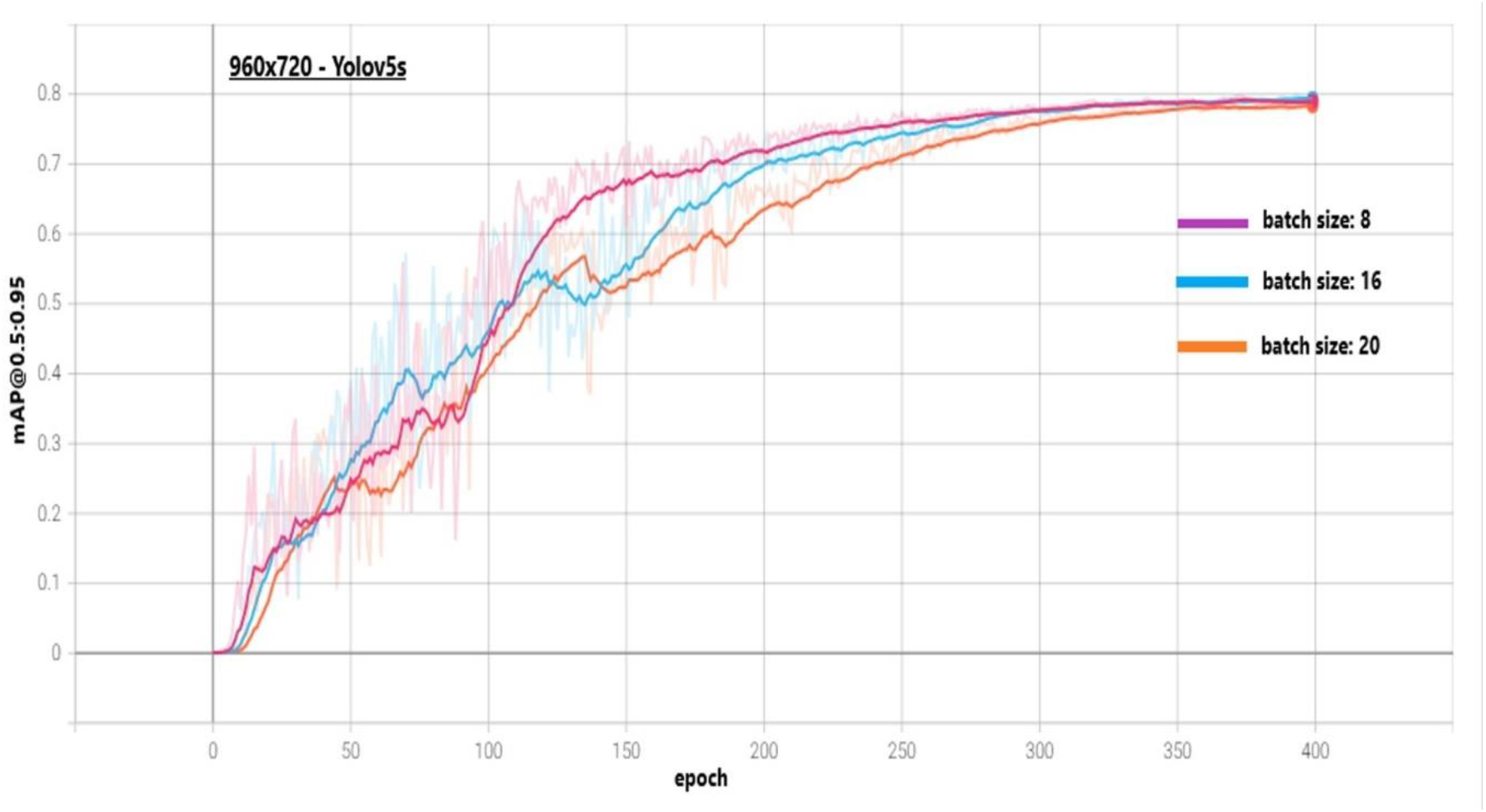
mAP@0.5:0.95 for different batch sizes

### Model Inference performance

The test set images representing varying scenarios were used for inference for all image size-batch size-model combinations. A summary of the inference results are presented in table 2.

**Table 2.** Inference results on test images

| Image Size | Batch size | YOLOv5n |  | YOLOv5s |  |
| --- | --- | --- | --- | --- | --- |
|  |  | Inference speed (ms/img) | Inference speed (fps) | Inference speed (ms/img) | Inference speed (fps) |
| 960x720 | 20 | 32.4 | 30.9 | 32.4 | 30.9 |
|  | 16 | 32.6 | 30.7 | 30.8 | 32.5 |
|  | 8 | 32.7 | 30.6 | 33.4 | 29.9 |
| 640x480 | 32 | 28.5 | 35.1 | 27.6 | 36.2 |
|  | 16 | 30 | 33.3 | 28.9 | 34.6 |
|  | 8 | 27.5 | 36.4 | 29.1 | 34.4 |
| 512x384 | 64 | 29.5 | 33.9 | 32.1 | 31.2 |
|  | 32 | 26.9 | 37.2 | 29.6 | 33.8 |
|  | 16 | 30.6 | 32.7 | 31.8 | 31.4 |
|  | 8 | 30.8 | 32.5 | 27.1 | 36.9 |
| 256x192 | 64 | 24.7 | 40.5 | 30.2 | 33.1 |
|  | 32 | 27.2 | 36.8 | 28.2 | 35.5 |
|  | 16 | 27.7 | 36.1 | 28.5 | 35.1 |
|  | 8 | 28.9 | 34.6 | 28.1 | 35.6 |

Fig. 8 shows a chart comparing the inference speeds using the best weights from all model training instances. The variation in inference speeds were only distinct for the highest and lowest image resolutions of 960×720 and 256×192 respectively; with the latter yielding inference speeds as low as 24.7millisecond per image (ms/img). Thus, the results do not clearly validate how variation in image resolution affects inference speeds. This could be explained by the fact that the inference was performed on a GPU that has more than enough memory and computational resources to process the images. Hence, we suppose that when the inferences are performed on highly memory-constrained processors like the NVIDIA Jetson GPUs typically used for mobile applications, the effect of the varying the input image sizes may be clearly seen.

**Fig. 8.**
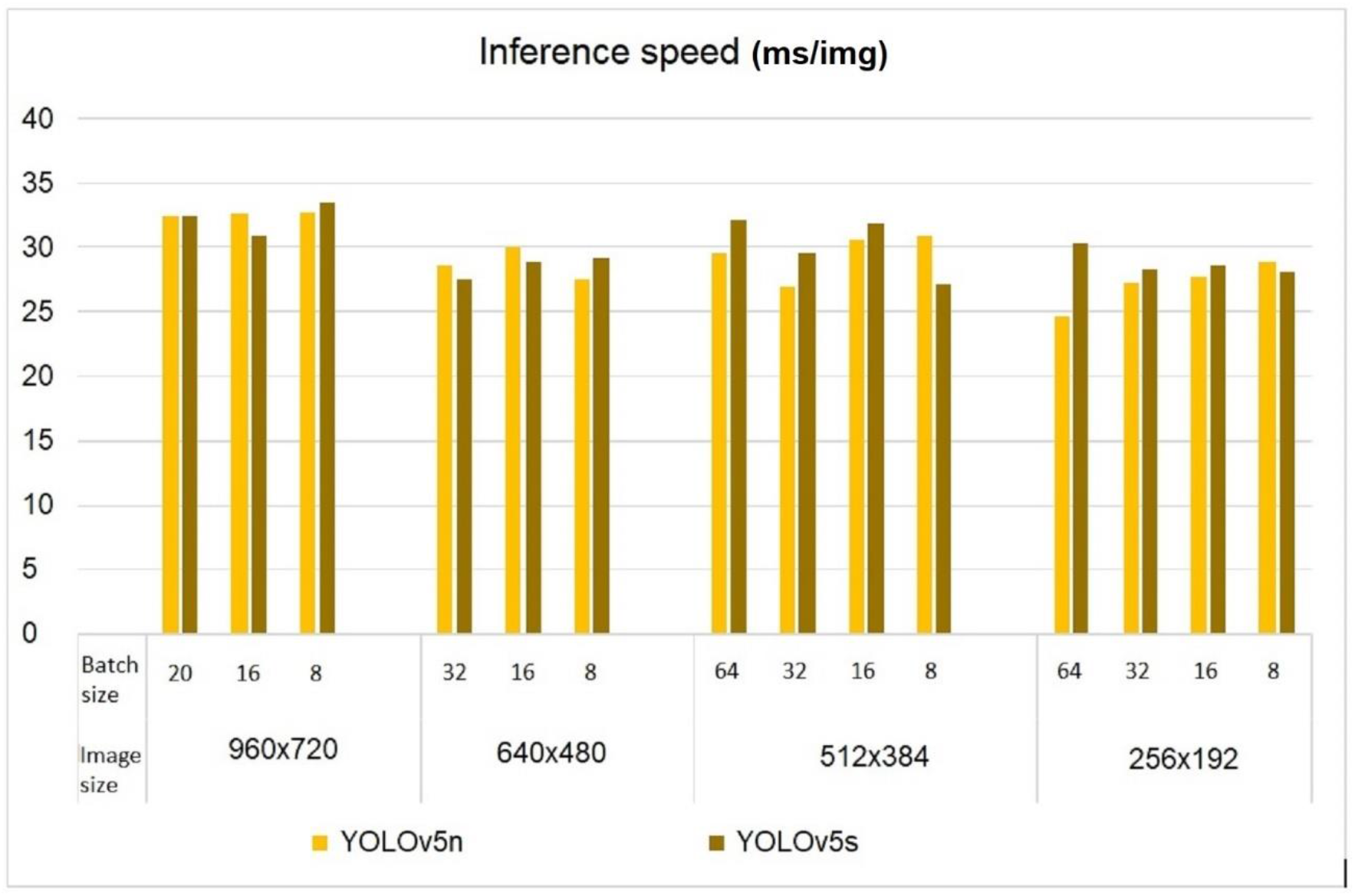
Inference speed of the models on test images

The longest inference time is 33.4ms/img, which corresponds to 29.9 fps, is more than sufficient for real-time applications. Thus, we conclude that for all image resolutions and training batch sizes of our dataset, the YOLOv5n and YOLOv5s models achieves inference speeds suitable for real-time applications.

#### Inference speed vs accuracy

Next, a plot of mAP@0.5:0.95 versus the inference speed in Fig. 9 was examined to highlight the potential speed-accuracy trade-offs. The choice of a model depends on the speed/accuracy requirements, which in turn depend on the application. The fastest model which is the YOLOv5n with 256×192 image resolution has the least mAP@0.5:0.95 at less than 0.35, whereas the model with best accuracy -YOLOv5s with 960×720 image resolution yields the slowest inference speed greater than 33ms/img.

**Fig. 9.**
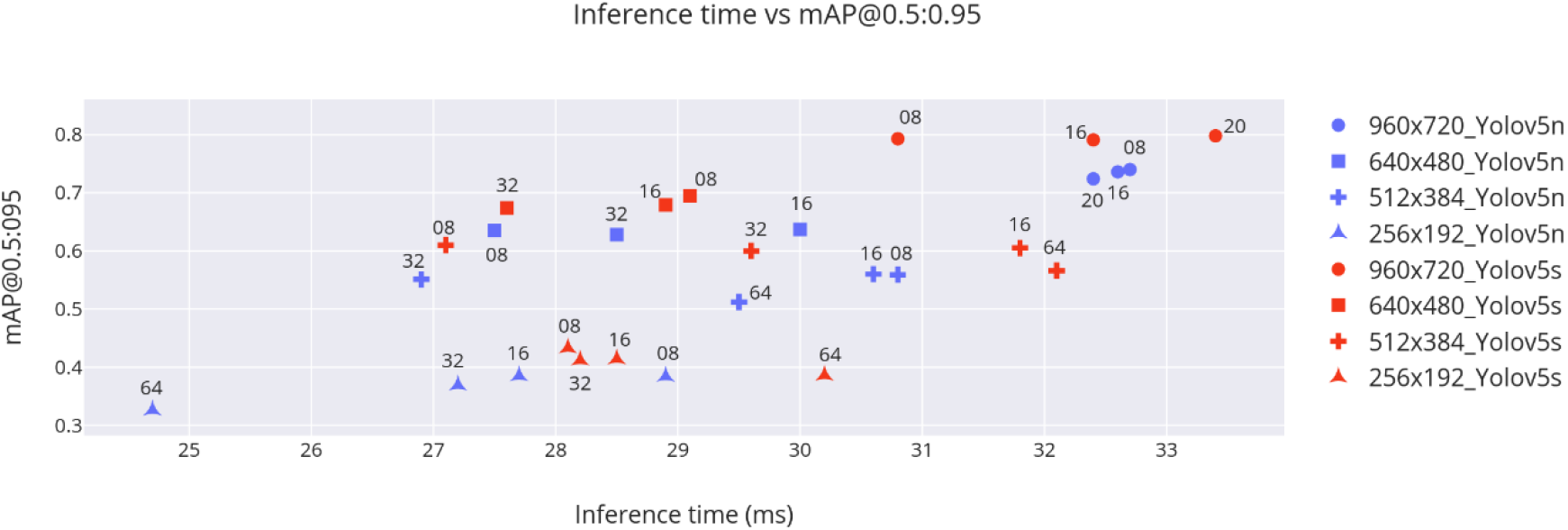
Plot of Inference time vs mAP@0.5:0.95 for the models

#### Performance of the inference on various field conditions

Since the inference speed of all the models were acceptable for real-time application, for comparison, a selection was done of the best performing model instance in terms of mAP@0.5:0.95 each for YOLOv5n and YOLOv5s. The first is the YOLOv5n with 960×720 input image resolution and training batch size of 8; the second is the YOLOv5s with 960×720 input image resolution and training batch size of 8. This selection of just two model instances was also done with the understanding that the results of the comparison across the models will be similar as you taper the input resolutions, but with the afore established attendant performance degradation that follows reduction in input image size.

##### Light conditions

Examination of the test images with even high lighting as well as with low lighting revealed that both model instances showed similar performance in terms of detecting the cassava objects. However, in high illumination images with large shadows from nearby trees, the YOLOv5n model missed some cassava objects under the shadow. This could be because the percentage of training images with large shadows was very little compared to images without shadows, therefore the model had insufficient images with shadows to learn from. This could be solved by collecting and re-training the model with more images with shadows. Moreover, this is not a major performance setback for real-life scenario given that large shadows (usually from tress) are a rare occurence in farms where UAVs are deployed for real-time weed control. Fig. 10 shows YOLOv5s and YOLOv5n models inferenced on the same image with shadow.

**Fig. 10.**
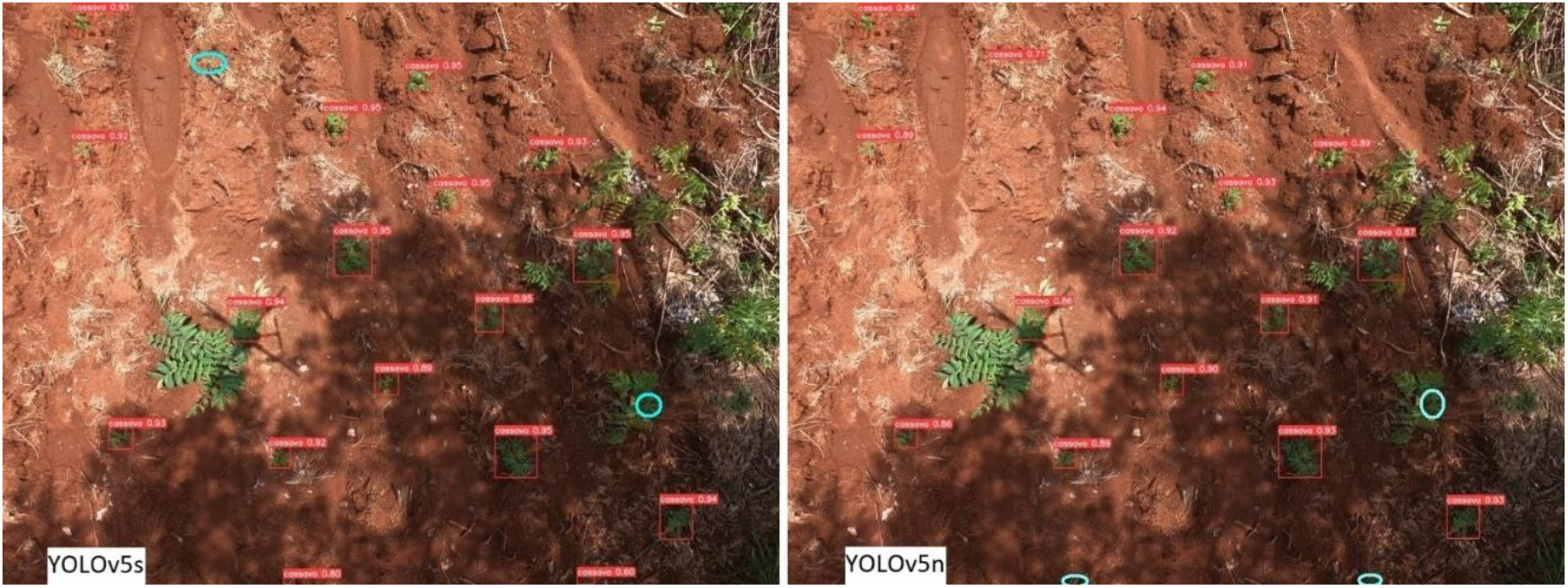
inference on images with shadow, missed cassava plants are circled

##### Growth stages

For images with early growth stage of the cassava, both model instances had no difficulty detecting almost all cassava objects. This high performance could be partly due to the fact that the early growth stage of the cassava corresponds to the stage of little or no weed on the farm. Therefore, the challenge of detecting cassava plants in the presence of leaf occlusions was absent. Both models also performed well on images of the later vegetative growth stage of the cassava. However, some cassava plants were missed by the YOLOv5n model. These were mainly the cassava plants that had delayed growth and still appeared little or were extremely occluded by the weeds. YOLOv5n model failed to detect a highly occluded cassava plant (circled in Fig. 11), which was successfully detected by the YOLOv5s model.

**Fig. 11.**
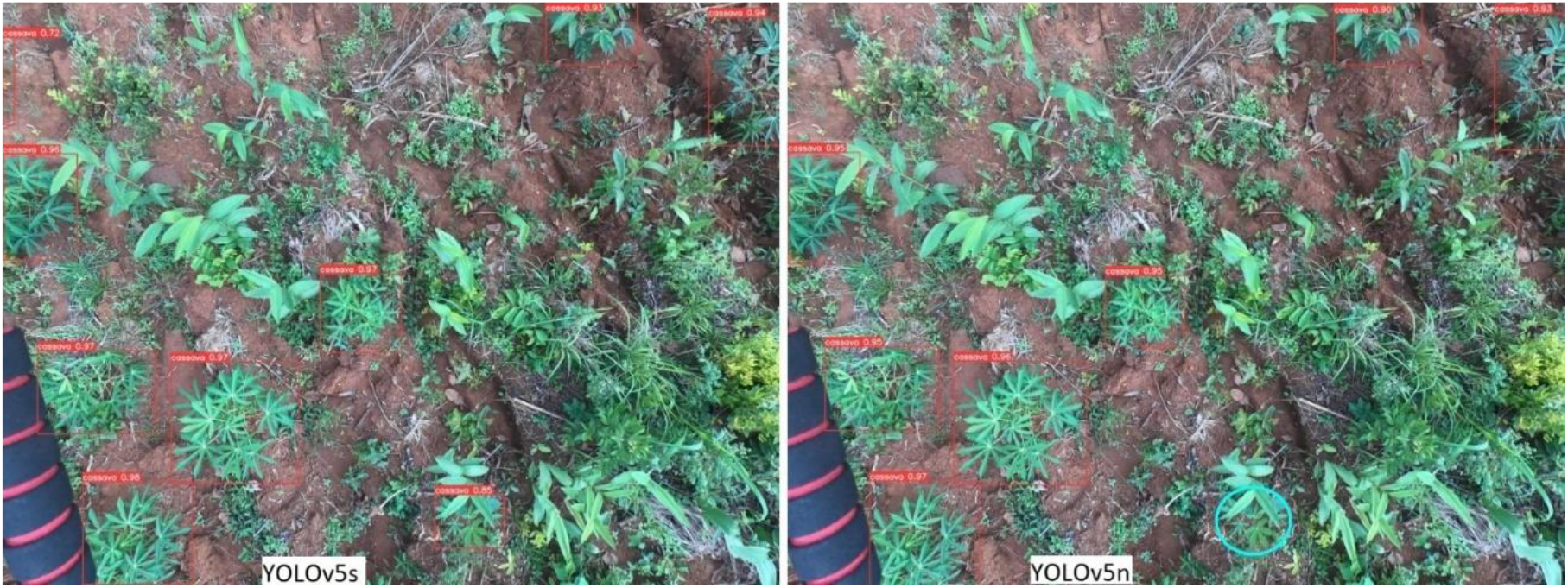
Inference on test image showing high leaf occlusion

##### Weed density

As the weed density increased, the degree of plant leaf occlusion increased. Though the models performed very well in detecting cassava plants on images of the early growth stage of cassava which corresponded to the period of almost zero weed density; however, for images with high weed density, it was observed that though the detection was good for both models, the YOLOv5n model predicted with less confidence than the YOLOv5s model for objects with high leaf occlusion. Also, some incorrect predictions with the YOLOv5n model were observed. Fig. 12 shows both models inferenced on the same test image with YOLOv5s yielding higher prediction confidence levels. The missed object is circled in Fig. 12(b).

**Fig. 12.**
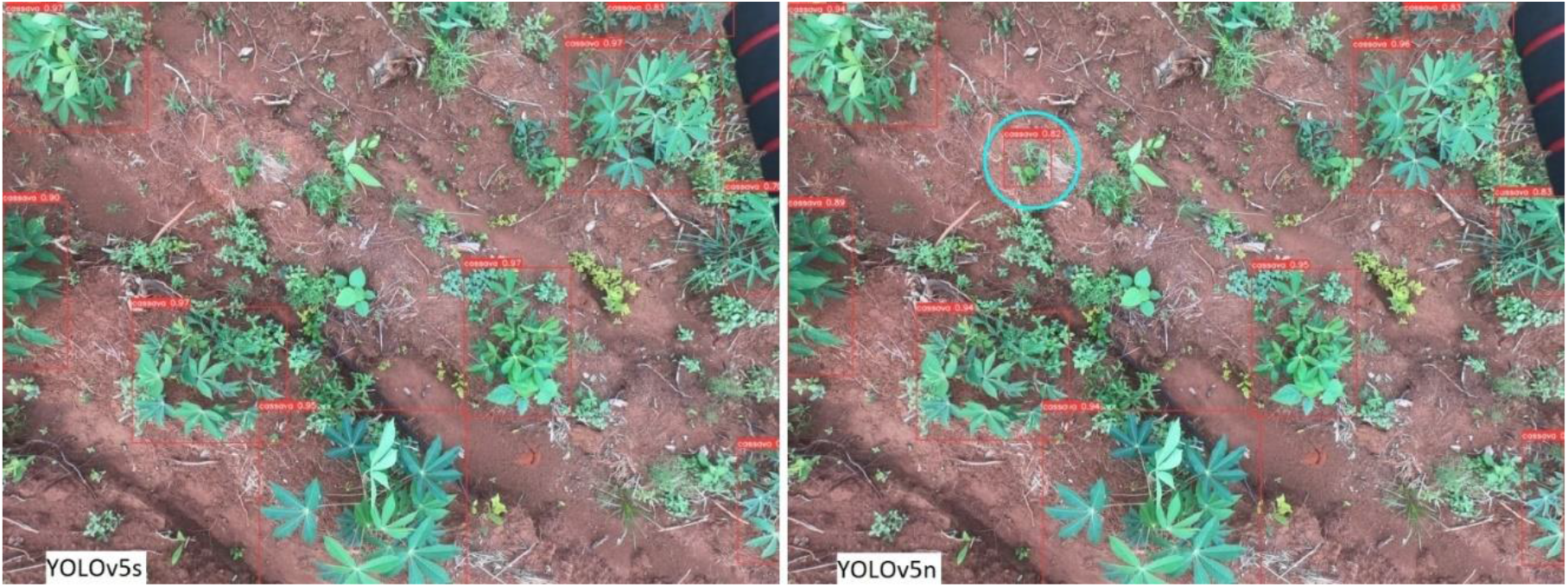
Inference image showing high weed density; YOLOv5s with generally higher prediction confidences, and YOLOv5n with generally lower prediction confidences

##### Row orientations

The inference results showed that both models performed well and there was no performance difference under varying row orientations. This good performance could be because of a few reasons. Firstly, the varying row orientations were fairly evenly distributed in the dataset, hence making the model learn better. Secondly, the image augmentation embedded in the models included image orientation. Thirdly, the structure of the cassava plant is radial and appears to be similar in shape despite the orientation of the rows, making the detections robust to row orientations.

## Conclusions

In this work, two YOLOv5 object detection models were evaluated for cassava plant detection for real-time applications. During model training, the image resolutions were varied, as well as the batch sizes. For both YOLOv5n and YOLOv5s, training time increases as the image resolution increases, but at the expense of decreasing accuracy. Generally, YOLOv5s showed higher accuracy than YOLOv5n for the same image resolution, but with greater training time and higher weight size. However, it was determined that training higher resolution images on YOLOv5n (which has smaller weights file) could yield better mAP@0.5:0.95 than training lower image resolutions on YOLOv5s, which is desirable for real-time applications where memory constraints are high. The speed vs accuracy plot highlights a range of possible speed/accuracy trade-offs to guide real-time deployment of the object detection model for cassava detection. The YOLOv5s slightly performed better than the YOLOv5n, however, the performance of both models under the various field conditions were shown to be impressive and, in our opinion, suitable for real-time cassava detection.

For future work, we intend to implement the models on a Jetson nano GPU mounted on a UAV and conduct field experiments to evaluate the model performances from field data. To further save computing power, the object detection model can be optimized to shrink the weights file size as well as reduce the complexity of the network so as to implement it on non-GPU mobile devices. We also plan to collect more data to completely cover all possible field conditions; thus, we can train a yet more robust model.

## Author Contributions

Conceptualization, E.C.N., O.A., O.I., and K.Y.; methodology, E.C.N., O.A., O.I., and K.Y; data curation, E.C.N.; writing—original draft preparation, E.C.N.; writing—review and editing, E.C.N., O.A., O.I., and K.Y.; supervision, O.A., O.I., and K.Y.; funding acquisition, E.C.N., O.A., O.I., and K.Y.; All authors have read and agreed to the submitted version of the manuscript.

## Data Availability Statement

The cassava dataset is available from the authors upon request. The data is not publicly available because the dataset is still being expanded and is a part of an ongoing funded project and will be published at a later date.

